# Blind Assessment of Monomeric AlphaFold2 Protein Structure Models with Experimental NMR Data

**DOI:** 10.1101/2023.01.22.525096

**Authors:** Ethan H. Li, Laura Spaman, Roberto Tejero, Yuanpeng Janet Huang, Theresa A. Ramelot, Keith J. Fraga, James H. Prestegard, Michael A. Kennedy, Gaetano T. Montelione

## Abstract

Recent advances in molecular modeling of protein structures are changing the field of structural biology. *AlphaFold-2* (AF2), an AI system developed by DeepMind, Inc., utilizes attention-based deep learning to predict models of protein structures with high accuracy relative to structures determined by X-ray crystallography and cryo-electron microscopy (cryoEM). Comparing AF2 models to structures determined using solution NMR data, both high similarities and distinct differences have been observed. Since AF2 was trained on X-ray crystal and cryoEM structures, we assessed how accurately AF2 can model small, monomeric, solution protein NMR structures which (i) were not used in the AF2 training data set, and (ii) did not have homologous structures in the Protein Data Bank at the time of AF2 training. We identified nine open source protein NMR data sets for such “blind” targets, including chemical shift, raw NMR FID data, NOESY peak lists, and (for 1 case) ^15^N-^1^H residual dipolar coupling data. For these nine small (70 - 108 residues) monomeric proteins, we generated AF2 prediction models and assessed how well these models fit to these experimental NMR data, using several well-established NMR structure validation tools. In most of these cases, the AF2 models fit the NMR data nearly as well, or sometimes better than, the corresponding NMR structure models previously deposited in the Protein Data Bank. These results provide benchmark NMR data for assessing new NMR data analysis and protein structure prediction methods. They also document the potential for using AF2 as a guiding tool in protein NMR data analysis, and more generally for hypothesis generation in structural biology research.

**Highlights:**

- AF2 models assessed against NMR data for 9 monomeric proteins not used in training.
- AF2 models fit NMR data almost as well as the experimentally-determined structures.
- *RPF-DP, PSVS*, and *PDBStat* software provide structure quality and RDC assessment.
- *RPF-DP* analysis using AF2 models suggests multiple conformational states.

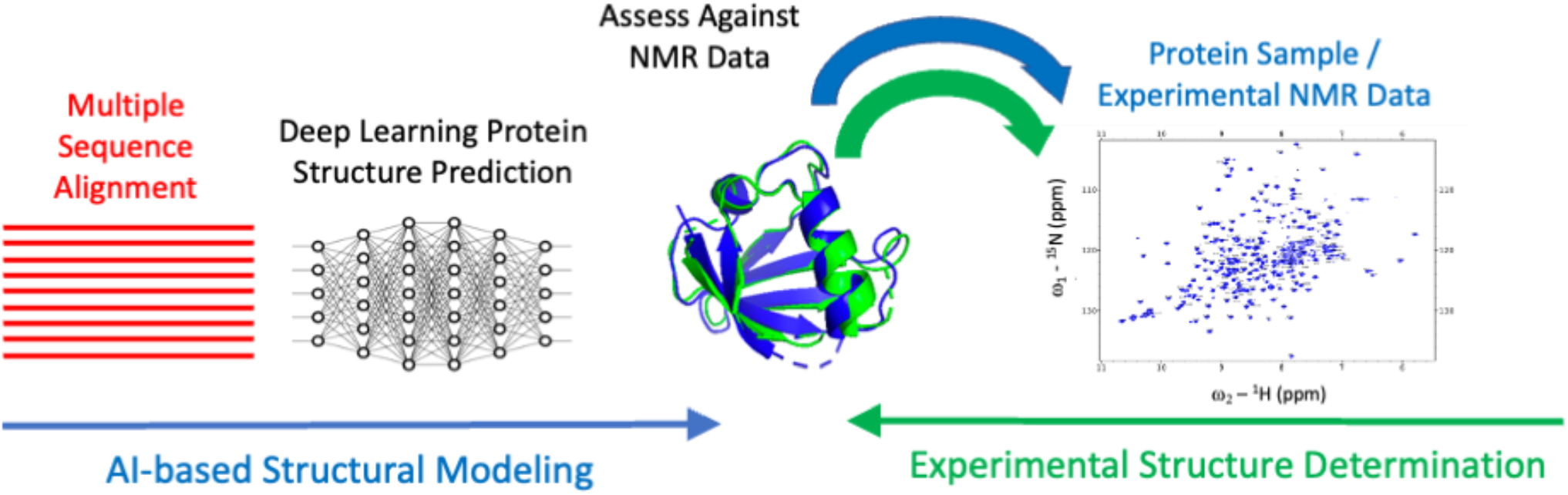

## 1. Introduction

Advances in protein structure modeling are having a high impact on the field of structural biology [1-5]. *AlphaFold-2* (AF2) [2, 6], an AI system developed by DeepMind, Inc, utilizes attention-based deep learning to predict models of protein structures, generally with high accuracy compared with structures determined experimentally by X-ray crystallography, NMR spectroscopy, and cryo-electron microscopy (cryoEM) [1, 7-9]. Advances in attention-based machine learning [10, 11] and contact prediction based on sequence covariance [12-17], along with the continuing rapid expansion of genomic sequence databases [18] and the growth of the open-access Protein Data Bank (PDB) [19], are enabling novel structural biology research.

These advances are driving the field of deep learning in structural biology. Along with other applications, AF2 models have the potential to revolutionize the processes used for NMR data and experimental biomolecular structure analysis, including methods for guiding NMR data analysis and experimental design.

In recent studies, we assessed models generated by AlphaFold against experimental NMR data [7, 8, 20, 21]. In most cases, AF2 models match the experimental Nuclear Overhauser effect (NOESY) peak list, chemical shift, and residual dipolar coupling (RDC) data nearly as well, or sometimes even better, than models deposited in the PDB based on experimental studies.

Special cases of discordance appear to result from the presence of multiple conformational states of the protein [8] or from errors in the experimental structure determination. In some cases, both NMR and X-ray crystal structures are available for the same protein construct, and comparisons with AF2, NMR, and X-ray crystal structure models against the corresponding NMR data are also largely consistent [7].

Although a few AF2 vs NMR data assessment studies have been carried out using proteins not included in the AF2 training data [8], in comparing AF2 models and NMR data for proteins available as both NMR and X-ray structure structures, one concern is the potential bias introduced if these X-ray crystal structures (or X-ray crystal structure of homologous proteins) were in the PDB at the time of the AF2 training, and were used in the AF2 training process.

Hence, the remarkable performance of AF2 on NMR / X-ray pairs [7] may have been impacted by its training data. As NMR structures were not used in the training of AF2 [2], these considerations pose the question of how accurately AF2 can model structures of proteins solved in solution by NMR methods, and for which structures of homologs were not in the PDB at the time of AF2 training.

Here, we test our hypothesis, building on our experiences in our previous studies [7, 8, 20, 21], that AF2 structural models are often nearly as accurate, or sometimes even more accurate, than published NMR-based structure models of small proteins, even where related protein crystal structures were not utilized in AF2 training. We collected open source NOESY peak list and FID data sets for protein NMR structures from the BioMagResDataBank (BMRB) [22] and recent literature [23]. We selected nine proteins for which the NOESY peak list data is available in standard *Sparky* [24] or *Xeasy* [25] format, and for which no X-ray crystal or cryoEM structures of homologs (using BLAST E_val < 10^−3^) are available in the PDB. For these nine proteins, we generated AF2 prediction models using the public Google Colab AF2-multimer server [2, 26], and evaluated these models against the corresponding NMR data (NOESY peak list and chemical shift data), residual dipolar coupling (RDC) data, and knowledge-based structure quality scores (e.g., backbone and sidechain dihedral angle distributions, *MolProbity* [27] packing scores, etc.), using the software packages Protein Structure Validation Server [28], *PDBStat* [29], and *RPF-DP* [30, 31]. In most of these cases, AF2 models have excellent knowledge-based structure quality scores and fit well to these NMR data, sometimes even better than the experimental structures deposited in the PDB. Overall, this study demonstrates the value of AlphaFold2 for modeling small monomeric protein structures, and supports its potential use in guiding analysis of experimental NMR data.

## 2. Methods

### Protein structure model validation

All structure quality statistical analyses were performed using the *Protein Structure Validation Software* (*PSVS*) suite version 2.0 [28] (https://montelionelab.chem.rpi.edu/PSVS/PSVS2/). *PSVS* runs a suite of software tools including *PDBStat* (ver 5.21.6), *ProCheck* (ver 3.5.4), *MolProbity* (mage ver 6.35.040409), *Cyrange* [32], and an implementation of the algorithms of *FindCore2* [33, 34] coded in *PDBStat. Procheck* and *MolProbity* structure validation scores were used to calculate a normalized Z score relative to the mean values and standard deviations of each of these scores obtained for 252 high-resolution X-ray crystal structures [28], with more positive Z scores indicating global model quality scores better than the average score (Z = 0) across this set of crystal structures.

### RPF-DP scores

RPF-DP scores are a set of fast and sensitive structure quality assessment measures that evaluate how well a 3D structure model fits with NOESY peak list and resonance assignment data. These protein NMR structure quality assessment metrics are described in detail elsewhere [8, 30, 31]. Briefly, Recall (R) measures the fraction of NOESY cross peaks that are consistent with short distances in query model structures, considering all possible assignments of each NOESY cross peak given the chemical shift list. Precision (P) measures the fraction of short proton pair distances in the query structure that are supported by a peak in the NOESY peak list, weighted by the interproton distance to minimize the impact of not observing weak NOEs arising from interproton distances near the edge of the defined distance cutoff [30]. The F-measure (F) is the harmonic mean of the Recall and Precision. The DP score is a scaled F-measure that accounts for lower-bound values of the F-measure, which would be expected for a random-coil structure (defines as DP = 0), and upper-bound values of F, which are related to the completeness of the NOESY data (defined as DP = 1). The global F and DP scores are types of “NMR R factors” that have been observed to be highly correlated with protein structure accuracy in several studies [7, 8, 20, 30, 31, 35].

Two methods were used for averaging DP scores across structural ensembles:

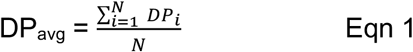

where DP_i_ is the DP score computed for the interproton distance matrix of the i-th conformer in the ensemble of N conformers, and

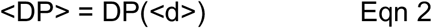

Where <d> is a distance matrix for which each element is the average interproton distance across the ensemble, and DP(<d>) is the DP score computed using <P>, <R>, and <F> determined using this distance matrix. In our experience, both of these averages are useful characterizations of how well the ensemble of conformers fit the NOESY data.

### Well-defined residue ranges

For the structure ensembles, the ranges of residues that are “well-defined”, were determined by standard conventions instantiated in the programs *Cyrange* [32] and *FindCore2* [33]. Residues were used in superimpositions and structure quality assessment only if they are both “well-defined” in the NMR and AlphaFold2 ensemble and “reliably predicted” (pLDDT > 80%) based on AlphaFold2 accuracy predictions.

### Residual dipolar coupling (RDC) analysis

^15^N-^1^H residual dipolar couplings, D_calc_, were calculated for individual model structures by single-value decomposition of the Saupe matrix [36] using *PDBStat* [29], called from *PSVS*, and plotted against the experimental values, D_exp_. Linear correlation coefficients r^2^ are also reported for these plots. Residual dipolar coupling quality (Q) scores were analyzed by *PDBStat* using both of the following two methods. The most commonly used RDC-fit score Q1, described by Cornilescu et al [37]) is

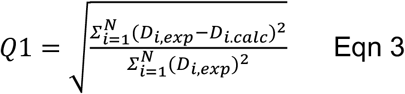

where D_exp_ and D_calc_ are the measured and calculated values of the RDC, and N is the number of RDCs assessed. In addition, we also assessed models using RDC-fit score Q2, described by Clore et al [38] and used by the DC: Servers for Dipolar Coupling Calculations (https://spin.niddk.nih.gov/bax/nmrserver/dc/svd.html).

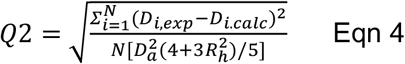

where D_a_ is the axial component, and R_h_ is the rhombic component of the orientation tensor. The Q2 factor is preferable in case of a limited RDC sampling over all possible orientations [38]. Regions of the protein structure with suspected flexibility, including not-well-defined regions and surface loops, were excluded from the RDC analysis.

### AlphaFold2 modeling

AlphaFold2 modeling was carried out on the Google ColabFold AF2-multimer server “AlphaFold2.ipynb” [2, 26]. This AI platform was trained using the PDB database of April, 2018 and did not use any NMR structures in the training data [2]. Modeling was done using default settings for generating multiple sequence alignments. No structure templates were used as input. The AF2 models were then relaxed with the amber_minimize.py protocol using OpenMM, which adds hydrogen atoms to the AF2 models, and reduces atomic clashes by minimizing conformational energy as defined by the *Amber99sb* [39] force field.

### Global Distance Test (GDT-TS) scores

GDT_TS scores were computed using backbone Cα atoms within the Comparison Residue Ranges, using the representative medoid model [29, 40] of the NMR model ensemble and the first-ranked AF2 model from the set of prediction models, by the method of Zemla [41], using the *TMScore* program [42] (downloaded from https://zhanggroup.org/TM-score/), as

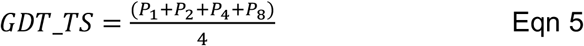

Here, P1, P2, P4, and P8 are the percent of residues with backbone Cα RMSD’s < 1 Å, < 2Å, < 4 Å, and < 8 Å, respectively, for consensus reliably modeled / well-define residue ranges of the superimposed structure pairs. GDT_TS = 100% would mean that all reliably-modeled residues superimpose with backbone Cα RMSD < 1 Å; while GDT_TS of 50% corresponds to an average backbone RMSD of about 4 Å. For brevity, GDT_TS scores are referred to throughout this paper as GDT scores, and are reported as real numbers between 0 (0%) and 1.0 (100%).

### RDC measurements

Samples for RDC measurements were prepared in 20 mM NH_4_OAc buffer, pH 4.5. containing 0.02 % NaN_3_, 10 mM DTT, 5 mM CaCL_2_ 100 mM NaCL, 10 % D_2_O, 50 μM DSS, 1 X Proteinase Inhibitor, at a protein concentration of 0.8 mM. ^15^N-^1^H RDC data were measured on samples partially aligned in polyacrylamide stretched gels (PAG) and polyethylene glycol (PEG) alignment media, as described previously [43].

### Superimpositions and molecular visualization

Molecular visualization for figures was done using *PyMol* [44]. The NMR and AF2 ensembles are shown using N, Cα, C’ backbone atoms. A darker color is used for well defined regions, with a lighter color for not-well-defined regions. Medoid conformers for NMR ensembles were identified, using *PDBStat*, as the conformer providing lowest RMSD to all other models in the ensemble [40]. The selected medoid was then trimmed to the appropriate comparison range from Table 2 and used as reference for superimposing the conformational ensemble.

**Table 1.** Nine Blind Protein NMR Targets.

| **Uniprot ID** | **NESG Target ID** | **PDB ID** | **DOI** | **BMRB ID** | **PDB Release Date** | **Residue (No of Residues)<sup>a</sup>** | **Authors** |
| --- | --- | --- | --- | --- | --- | --- | --- |
| O31898 | SR399 | 2HEQ | 10.2210/pd<br>b2HEQ/pdb | 7155 | 2006-08-15 | 2-71<br>(70) | Ramelot, T.A., Cort, J.R., Wang, D., Janjua, H., Cunningham, K., Ma, L.-C., Xiao, R., Liu, J., Baran, M., Swapna, G.V.T., Acton, T.B., Rost, B., Montelione, G.T., Kennedy, M.A. |
| Q8Q045 | MaR214A | 2KBN | 10.2210/pd<br>b2KBN/pdb | 16051 | 2008-12-03 | 67-166<br>(100) | Ramelot, T.A., Ding, K., Magliqui, M., Jiang, M., Ciccocanti, C., Xiao, R., Lui, J., Everett, J.K., Swapna, G., Acton, T.B., Rost, B., Montelione, G.T., Kennedy, M.A. |
| Q9KL30 | VcR52 | 2KHD | 10.2210/pd<br>b2KHD/pdb | 16238 | 2009-06-16 | 1-100<br>(100) | Ramelot, T.A., Cort, J.R., Wang, H., Ciccocanti, C., Jiang, M., Liu, J., Rost, B., Swapna, G.V.T., Acton, T.B., Xiao, R., Everett, J.K., Montelione, G.T., Kennedy, M.A. |
| Q4L7R3 | ShR105F | 2KIW | 10.2210/pd<br>b2KIW/pdb | 16298 | 2009-07-07 | 64-166<br>(103) | Yang, Y., Ramelot, T.A., Belote, R.L., Foote, E.L., Janjua, H., Nair, R., Rost, B., Swapna, G., Acton, T.B., Xiao, R., Everett, J.K., Montelione, G.T., Kennedy, M.A. |
| A1ZBM2 | FR629A | 2KJR | 10.2210/pd<br>b2KJR/pdb | 16338 | 2009-06-23 | 8-92<br>(85) | Ramelot, T.A., Cort, J.R., Shastry, R., Ciccocanti, C., Jiang, M., Nair, R., Rost, B., Swapna, G., Acton, T.B., Xiao, R., Everett, J.K., Montelione, G.T., Kennedy, M.A. |
| A7VTE5 | QIR8A | 2KOB | 10.2210/pd<br>b2KOB/pdb | 16498 | 2009-09-26 | 61-160<br>(100) | Ramelot, T.A., Lee, D., Ciccocanti, C., Jiang, M., Nair, R., Rost, B., Acton, T.B., Xiao, R., Everett, J.K., Montelione, G.T., Kennedy, M.A. |
| Q7P141 | CvR118A | 2KZV | 10.2210/pd<br>b2KZV/pdb | 17020 | 2010-08-25 | 175-257<br>(83) | Yang, Y., Ramelot, T.A., Wang, D., Ciccocanti, C., Mao, L., Janjua, H., Acton, T.B., Xiao, R., Everett, J.K., Montelione, G.T., Kennedy, M.A. |
| Q5XPI4 | HR8700A | 2MA6 | 10.2210/pd<br>b2MA6/pdb | 19329 | 2013-07-31 | 1247-1304<br>(58) | Ramelot, T.A., Yang, Y., Janjua, H., Kohan, E., Wang, H., Xiao, R., Acton, T.B., Everett, J.K., Montelione, G.T., Kennedy, M.A. |
| A0A0H2<br>VTH7 | SfR125 | 2RN7 | 10.2210/pd<br>b2RN7/pdb | 11017 | 2008-01-15 | 1-108<br>(108) | Ramelot, T.A., Cort, J.R., Semesi, A., Garcia, M., Yee, A.A., Arrowsmith, C.H., Kennedy, M.A. |

<sup>a</sup>Uniprot residue ranges and number of residues in deposited NMR structure, excluding disordered purification tags, if any.

**Table 2.** Well-defined and Comparison Residue Ranges for NMR and AF2 model ensembles.

| PDB ID | Method | Consensus Well-defined Residue Range | Comparison Residue Range<sup>a</sup> | GDT Score for Comparison Residue Range | Backbone (N,Cα,C') rmsd for Comparison Residue Range<sup>d</sup> | GDT Score for Full Length Protein<sup>d</sup> |
| --- | --- | --- | --- | --- | --- | --- |
|  |  |  |  | AF2 Model 1<sup>b</sup><br>(AF2 Best) | AF2 Model 1<sup>b</sup><br>(AF2 Best) | AF2 Model 1<sup>b</sup><br>(AF2 Best) |
| 2HEQ | NMR | 11-20, 33-41, 46-68 | 14-20, 33-41, 47-68 | 0.97<br>( 0.97 ) | 0.65 Å<br>( 0.65 Å ) | 0.68<br>( 0.69 ) |
|  | AF2 | 14-23, 32-42, 47-69 |  |  |  |  |
| 2KBN | NMR | 3-14, 17-29, 35-53, 58-77, 81-86, 89-97 | 5-14, 17-28, 35-53, 58-77, 81-86, 89-97 | 0.95<br>(0.96) | 0.85 Å<br>(0.79 Å) | 0.88<br>(0.88) |
|  | AF2 | 5-28, 33-100 |  |  |  |  |
| 2KHD | NMR | 31-38, 40-58, 62-69, 75-81, 84-97 | 31-38, 42-58, 62-66, 75-81, 84-95 | 0.93<br>(0.93) | 0.99 Å<br>(0.99 Å) | 0.63<br>(0.66) |
|  | AF2 | 30-95 |  |  |  |  |
| 2KIW | NMR | 5-17, 21-87 | 5-15, 22-36, 38-87 | 0.81<br>(0.81) | 1.90 Å<br>(1.90 Å) | 0.66<br>(0.66) |
|  | AF2 | 2-15, 22-36, 38-90 |  |  |  |  |
| 2KJR | NMR | 15-22, 27-52,<br>55-82, 85-94 | 15-22, 32-52, 55-82,<br>85-94 | 0.92<br>(0.94) | 1.08 Å<br>(0.97 Å) | 0.79<br>(0.80) |
|  | AF2 | 15-23, 32-94 |  |  |  |  |
| 2KOB | NMR | 63-157 | 63-156 | 0.97<br>(0.98) | 0.75 Å<br>(0.69 Å) | 0.87<br>(0.88) |
|  | AF2 | 63-156 |  |  |  |  |
| 2KZV | NMR | 8-21, 27-39,<br>45, 53-59.<br>73-80 | 14-20, 27-38, 57-59,<br>73-80 | 0.94<br>(0.96) | 1.03 Å<br>(0.90 Å) | 0.66<br>(0.68) |
|  | AF2 | 14-20, 26-38, 57-80 |  |  |  |  |
| 2MA6 | NMR | 10-17, 20-44,<br>49-58 | 10-17, 20-44, 49-58 | 0.97<br>(0.97) | 0.84 Å<br>(0.83 Å) | 0.88<br>(0.88) |
|  | AF2 | 9-59 |  |  |  |  |
| 2RN7 | NMR | 8-25, 27-55 | 8-25, 27-54 | 0.95<br>(0.96) | 0.81 Å<br>(0.79 Å) | 0.49<br>(0.50) |
|  | AF2 | 8-54 |  |  |  |  |

<sup>a</sup>The Comparison Residue Range is the consensus between NMR and AF Consensus Residue Ranges.
<sup>b</sup>Comparing the medoid of the NMR ensemble with “Model 1”, defined as the conformer ranked number 1 of the AF2 ensemble (and with the “AF2 Best” model, defined as the model with the highest GDT score with the medoid NMR structure)

### Software accessibility

All software used in this study and developed by the Montelione laboratory are available under open source licenses in a public GitHub site: https://github.rpi.edu/RPIBioinformatics or as on-line servers. Software packages developed by third-parties are publicly available from the url’s provided here and/or in the original references cited.

## 3. Results

Experimentally-determined protein structure coordinates and NMR data were initially considered for 100 small, monomeric proteins utilized in a recent study using *ARTINA*, an artificial intelligence and *CYANA-*based program for automated peak picking, resonance assignment, and structural modeling [23]. Most of these protein NMR structures and data were deposited in the public PDB by the Northeast Structural Genomics Consortium (NESG) [45], and the corresponding NMR data was downloaded from the BMRB.

Relatively few NMR structure depositions include NOESY FID and peak list data needed for the model assessment methods utilized in this study. This is because such raw NMR data deposition is not required or fully supported by the BMRB and PDB. Here, we focused on small (< 110 residues) monomeric protein NMR structures that (i) have not also been determined using X-ray crystallography or cryoEM, (ii) have publicly-available NOESY FID and NOESY peak list data, and (iii) were not used in the AF2 training [2]. This selection process is outlined in Figure 1.

**Figure 1.**
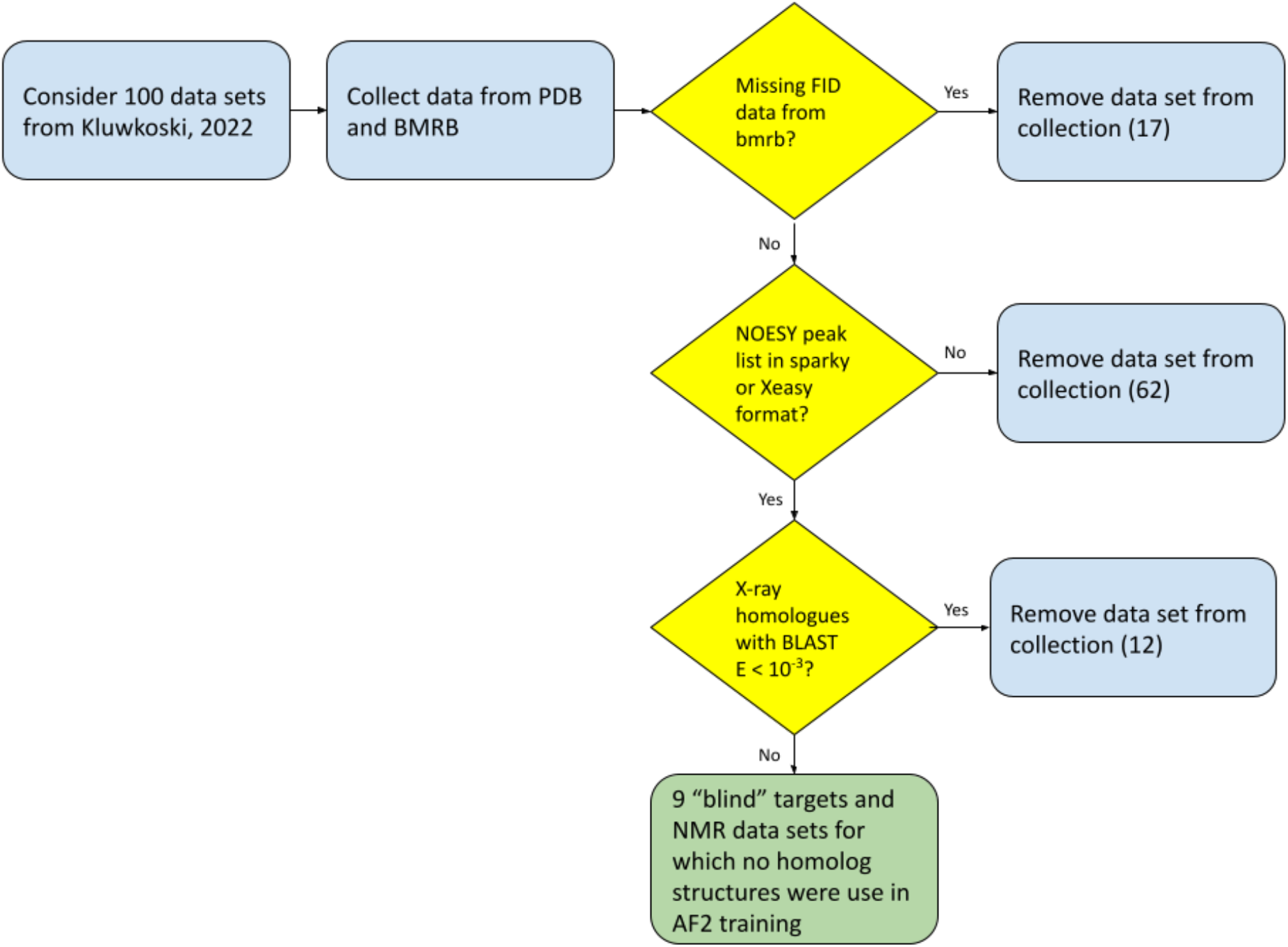
Target Selection Flowchart. These nine “blind” protein NMR targets are summarized in Table 1, along with their Uniprot ID, NESG Target ID, PDB ID, PDB entry DOI, BMRB ID, PDB Release Date, Residue Range (and number of residues), and PDB entry author list. Of these, only 1 (Uniprot ID Q7P141 (residues 175-257); PDB ID 2KZV) has ^15^N-^1^H residual dipolar coupling (RDC) data (for two orientation media) deposited in the PDB that are suitable for this assessment study. Interestingly, due to the nature of the data sets organized by Kluwkoski, et al. [23], which ideally included NOESY and triple-resonance NMR FID data sets together with spectral processing scripts, the final set of 9 data sets were all from the NESG research program. Aside from deposition in the PDB, none of these NMR structures have been previously published. NMR data (including raw FID data), NMR-based structure models, and AF2 structure prediction models for these nine protein targets, together with key input and output files and analysis results needed to reproduce this study, were then organized in a public GitHub Site to provide a resource for future studies (Supplementary Material).

We began by considering the 100 protein NMR data sets collated by Kluwkoski et al [23]. First, seventeen (17) data sets lacking NOESY FID data were removed from this collection. Next, sixty-two (62) protein targets for which NOESY FID data is available, but the NOESY peak list data was not deposited by the spectroscopist, were also culled. As these targets do have NOESY FID data, it would be possible to process and peak pick these NOESY spectra for future studies; however in this study we wanted to use exactly the same NOESY peak lists used by the spectroscopist in their final experimental structure analysis as deposited in the BMRB. Lastly, a filtered search for homologs on known 3D structure utilizing the *Gapped BLAST* tool integrated in the RCSB PDB server (https://www.rcsb.org/) was executed (on July 25, 2022).

Using an E-value of < 10^−3^, a generous cutoff for removing remote homologs of known structure, twelve (12) of the remaining twenty-one (21) targets were found to have 3D structures in the PDB that may have been used in training of AF2, and were removed from the study target set. The resulting nine (9) monomeric protein targets and NMR data sets were then identified as “blind” targets, not used in AF2 training, suitable for this study.

In comparing atomic coordinates for experimental and predicted protein structures, it is critical to account for the fact that some parts of the protein structure are not reliably modeled by the experimental method, the prediction method, or by both methods. Failure to account for the “not-well-defined” regions of these structures can result in erroneous conclusions, as the superimposition can be dominated by the diverse conformations in these regions of unreliably-modeled structure, resulting in poor superimposition of the well-defined regions. For the experimental NMR ensembles, well-defined vs. not-well-defined structure regions were determined by the consensus of three commonly used methods and conventions: the backbone dihedral angle order parameter analysis introduced by Hyberts and Wagner [46], the Cyrange method developed by Kirchner and Güntert [32], and the FindCore2 method developed by Snyder, Montelione and co-workers [33, 34], each implemented in the *PSVS* ver 2.0 server.

The well-defined residue ranges reported for each of these methods are summarized in Supplementary Table S1 for NMR structure ensembles of the 9 targets, along with consensus well-defined residue ranges. Generally, the three methods are in very good agreement. For AF2 models, results of the same “well defined regions” analysis, using the top 5 conformers generated by the AF2 server, are also reported in Supplementary Table S1, along with residue ranges predicted by AF2 to have “high modeling confidence” (i.e. pLDDT > 80) [2]. For these AF2 models, the well-defined residue ranges are also consistent across these four methods, and are also reported in Supplementary Table S1, together with corresponding Consensus Residue Ranges. Comparing the consensus well-defined ranges for the NMR and AF2 ensembles allows identification of structure Comparison Residue Ranges; i.e. the subset of residues for which modeling is judged to be “well defined” in both the NMR and AF2 ensembles (Table 2). These Comparison Residue Ranges were then used for structure superimposition and computation of GDT scores and backbone root-mean-square-deviations (RMSDs), also summarized in Table 2. As a standard convention, we used the medoid conformation of the NMR ensemble (conformer most like the other conformers) and the top ranked (AF2 Model 1) conformer of the AF2 ensemble (most confident prediction) to compute GDT scores comparing the NMR and AF2 models. Additionally, a full length GDT_TS score, including all residues in the superimposition, is reported to demonstrate the significant impact of including not-well defined regions in these structural comparisons.

Considering the Comparison Residue Ranges, excluding one outlier (PDB ID 2KIW), the NMR (medoid) and AF2 (rank 1) models are in excellent agreement, with GDT scores (backbone rmsd) ranging from 0.92 - 0.97 (0.65 - 1.08 Å), and average GDT score for 8 targets of 0.95 (and backbone rmsd 0.88 Å) (Table 2). These structural differences between the NMR and AF2 models are within the experimental uncertainty of the NMR models. The one outlier in Table 2 is target PDB_ID 2KIW, with GDT = 0.81 (backbone rmsd = 1.90 Å). These structural superimpositions are shown graphically in Figure 2. Note that considering also the not-well-defined regions of these 8 targets results in much poorer superimpositions (e.g. GDTs ranging from 0.49 - 0.88, Table 2), as these inaccurate regions of the NMR and/or AF2 models negatively impact these structural superimpositions. This illustrates the importance of using well-defined residues when assessing NMR structures against prediction models.

**Figure 2.**
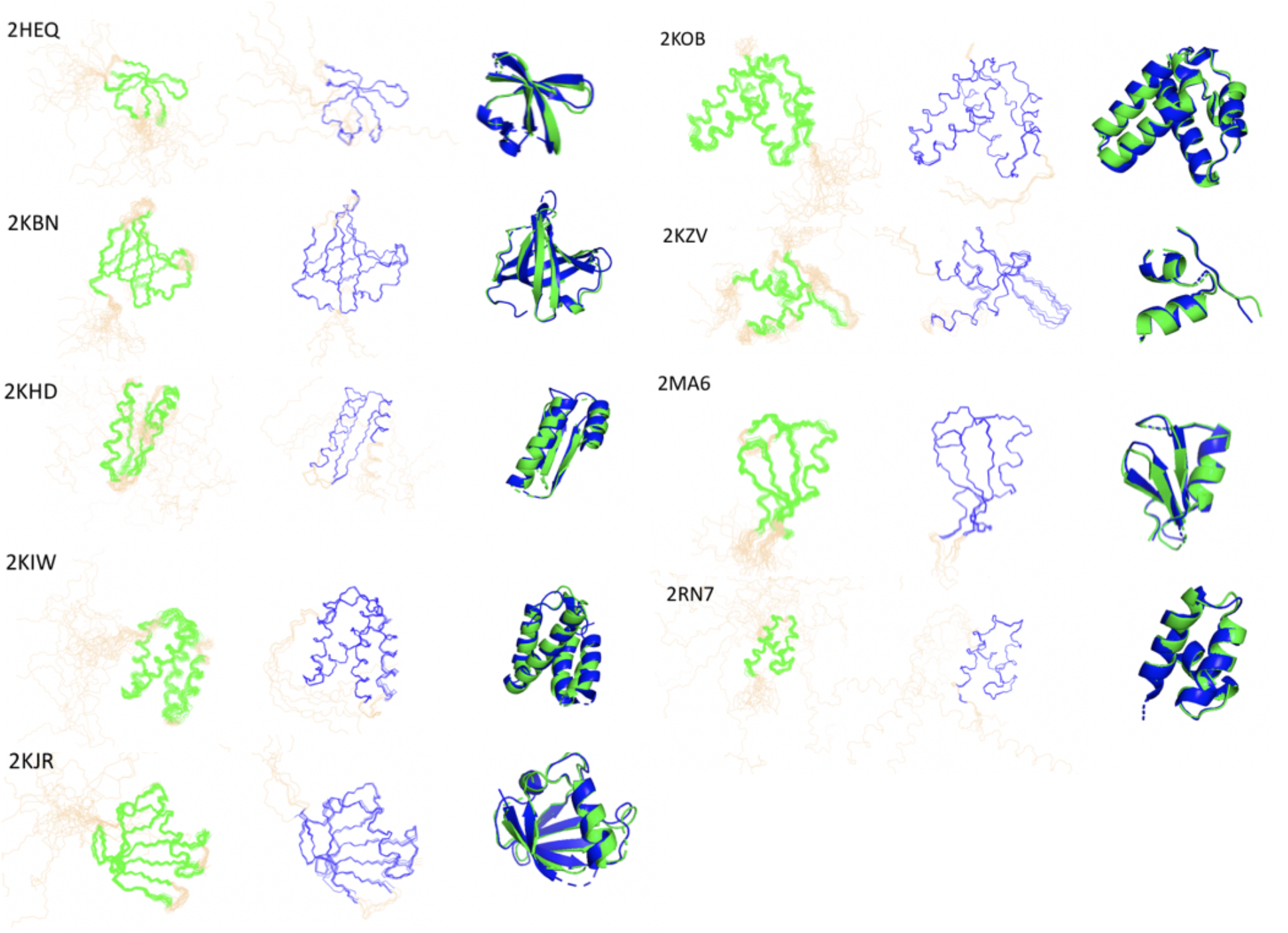
NMR and AF2 ensemble backbone structure comparisons. For each target: left – NMR ensemble [green]; center – AF2 ensemble [blue]; right – NMR medoid conformer [green] superimposed on AF2 Model 1 conformer (blue), for Residue Comparison Range superimposition. The not-well-defined regions of the NMR or AF2 ensembles are shown in light brown in the ensemble representations. Only residues in the Comparison Residue Ranges are shown in the ribbon representations.

In order to assess the energetics of both backbone and sidechains in these NMR and AF2 models, the structures were also analyzed using the metrics of *ProCheck-bb* (backbone dihedral angle distributions) [47], *ProCheck-all* (backbone and sidechain dihedral angle distributions) [47], *MolProbity* atom packing scores [27], and Richardson backbone Ramachandran statistics [48], as implemented in the *PSVS* server [28]. These results, reported as Z-scores relative to high-resolution (< 1.8 Å) X-ray crystal structures [28], are summarized in Table 3. More positive Z scores are more energetically-favorable. Generally, Z scores > -3 are typical of good quality NMR structures. While all of the NMR and AF2 models satisfy this criterion, the AF2 models generally exhibit better *MolProbity* packing scores than the NMR structures, with values typical of very good quality core sidechain structures. For some targets, *ProCheck-bb* and *ProCheck-all* scores are better for the NMR models, and for other targets they are better for the AF2 models. In most cases, these structure quality Z scores are either better, or no more than one standard deviation (ΔZ = -1) poorer, for the AF2 structures compared with the corresponding NMR structure. The Richardson Ramachandran statistics (Table 3) are generally better for the NMR models.

**Table 3.** Protein Structure Validation Software Suite (PSVS) scores for 9 protein targets.

| Protein | Method | ProCheck -<br>bb<sup>a</sup> | ProCheck -<br>all<sup>a</sup> | MolProbity<sup>a</sup> | Ramachandran<br>Statistics<sup>b</sup><br>(percent most favored /<br>allowed / disallowed) |
| --- | --- | --- | --- | --- | --- |
| 2HEQ | NMR | **-1.49<sup>c</sup>** | -2.07 | -1.73 | **99.5 / 0.2 / 0.2** |
|  | AF2 | -1.81 | **-1.83** | **0.63** | 93.7 / 5.1 / 1.2 |
| 2KBN | NMR | **-1.46** | -1.42 | -1.83 | **95.5 / 4.4 / 0.1** |
|  | AF2 | -1.57 | **-0.95** | **1.53** | **94.2 / 5.3 / 0.6** |
| 2KHD | NMR | **1.85** | **1.66** | -2.00 | **99.3 / 0.7 / 0.0** |
|  | AF2 | -0.55 | -0.59 | **1.53** | 90.8 / 8.6 / 0.6 |
| 2KIW | NMR | **2.79** | **2.72** | -1.47 | **98.8 / 1.0 / 0.1** |
|  | AF2 | 1.34 | 1.95 | **1.19** | 95.4 / 4.2 / 0.4 |
| 2KJR | NMR | -0.31 | **0.30** | -0.91 | 94.2 / 5.1 / 0.7 |
|  | AF2 | **-0.16** | 0.18 | **1.53** | **98.2 / 1.8 / 0.0** |
| 2KOB | NMR | 1.38 | 0.95 | -1.71 | 98.5 / 1.2 / 0.3 |
|  | AF2 | **1.61** | **1.60** | **1.53** | **99.4 / 0.6 / 0.0** |
| 2KZV | NMR | **1.10** | **1.18** | -1.37 | **97.9 / 2.0 / 0.2** |
|  | AF2 | -0.16 | 0.06 | **1.25** | 95.6 / 2.6 / 1.8 |
| 2MA6 | NMR | -2.91 | -2.78 | -0.20 | 86.4 / 13.4 / 0.2 |
|  | AF2 | **-2.08** | **-2.19** | > 3.00<sup>d</sup> | **92.3 / 6.0 / 1.8** |
| 2RN7 | NMR | **1.61** | **1.18** | -0.59 | **98.4 / 1.5 / 0.1** |
|  | AF2 | 1.34 | **1.18** | **1.53** | 95.3 / 4.5 / 0.2 |

<sup>a</sup>Z scores relative to the corresponding scores obtained from a database of 252 high-resolution X-ray crystal structures solved at < 1.8 Å resolution (Bhattacharya et al., 2007). <sup>b</sup>Percent of backbone residues in percent most favored / allowed / disallowed of Ramachandran map (Lovell et al., 2003). <sup>c</sup> For each NMR / AF2 comparison, and for each metric, the better score (more positive Z-score) is indicated in bold font. <sup>d</sup> When zero *Molprobability* class scores are present, the z score is set to > 3.00.

In addition to the knowledge-based structural quality scores summarized in Table 3, the *PDBStat* software was used to check for other irregular structure quality issues in the NMR and AF2 models The scripts used for this analysis are presented in Supplementary Material.Unusual trans peptide bond dihedral angles ω, defined as values outside the range of 180 ± 40 deg, were observed in several AF2 models, but not in any of the NMR structure models. For target PDB ID 2MAF, all five AF2 models have cis peptide bonds at Gln24-Pro25, which is consistent with the NMR data. For target PDB ID 2KZV, one AF2 model (ranked 2^ed^) has a non-proline cis peptide bond at residue His89, which is not consistent with the NMR data. None of the NMR or AF2 models have mirror image D-amino acid residues.

*RPF-DP* scores, comparing models with chemical shift and NOESY peak list data, were calculated in the *RPF* web server (https://montelionelab.chem.rpi.edu/rpf/) [31], using the chemical shift list downloaded from the BMRB, the PDB coordinate file, and the NOESY peak list data provided with the PDB deposition. *RPF-DP* scores compare interproton distances in a model against NOESY peak list and chemical shift data, and are sensitive to the accuracy of both the backbone and the buried sidechains of the protein structure. For each method, DP_min_ (the lowest DP score across all of the models), DP_max_ (the highest DP score across all of the models), DP_avg_, <DP>, <R>, <P>, and <F> were calculated. Generally, good NMR structures have DP_avg_ > 0.60 and <DP> > 0.70 [49], although for very good NOESY data sets and accurate, relatively-rigid structures these values can be > 0.90. These results are summarized in Table 4. The AF2 structures have DP_avg_ ranging from 0.61 to 0.78, and <DP> ranging from 0.65 to 0.80. These ranges are similar to those of the NMR structures (0.57 - 0.77 and 0.71- 0.84, respectively). For the DP_min_, DP_max_, and DP_avg_ metrics, the AF2 models generally have better scores than the NMR models, while for <DP> metric the experimental NMR models generally provide somewhat better scores. Even though the NMR structures are modeled using distance restraints based on these NOESY data, the AF2 models often fit these NMR data as well, or sometimes even better, than the experimental NMR structures deposited in the PDB. In some cases, these DP scores are lower for the AF2 models than the experimental NMR structures, indicating that more accurate modeling (i.e. models that better fit the data) is possible.

**Table 4.** RPF Scores for NMR and AF2 model ensembles.

| Protein | Method | DP<sub>min</sub><sup>a</sup> | DP<sub>max</sub><sup>b,c</sup> | DP<sub>avg</sub> | <DP><sup>d</sup> | <R> | <P> | <F> |
| --- | --- | --- | --- | --- | --- | --- | --- | --- |
| 2HEQ | NMR | 0.68 | 0.74 | 0.71 | **0.78<sup>d</sup>** | **0.89** | **0.94** | **0.91** |
|  | AF2 | **0.74** | **0.78** | **0.77** | **0.78** | **0.89** | 0.93 | **0.91** |
| 2KBN | NMR | 0.67 | 0.72 | 0.70 | **0.78** | 0.92 | **0.81** | **0.86** |
|  | AF2 | **0.74** | **0.76** | **0.75** | 0.77 | **0.93** | 0.80 | 0.86 |
| 2KHD | NMR | 0.55 | 0.64 | 0.59 | **0.71** | **0.96** | **0.86** | **0.91** |
|  | AF2 | **0.61** | **0.65** | **0.63** | 0.65 | **0.96** | 0.84 | 0.90 |
| 2KIW | NMR | 0.50 | 0.61 | 0.57 | **0.74** | 0.95 | **0.80** | **0.87** |
|  | AF2 | **0.59** | **0.63** | **0.61** | 0.65 | **0.97** | 0.77 | 0.86 |
| 2KJR | NMR | **0.74** | 0.79 | **0.77** | **0.83** | **0.97** | **0.90** | **0.93** |
|  | AF2 | **0.74** | **0.80** | 0.76 | 0.79 | **0.97** | 0.88 | **0.93** |
| 2KOB | NMR | 0.73 | **0.79** | 0.76 | **0.84** | **0.96** | **0.86** | **0.91** |
|  | AF2 | **0.75** | 0.78 | **0.78** | 0.78 | **0.96** | 0.84 | 0.90 |
| 2KZV | NMR | 0.63 | **0.71** | 0.68 | **0.82** | **0.98** | **0.91** | **0.94** |
|  | AF2 | **0.68** | **0.71** | **0.70** | 0.73 | 0.97 | 0.88 | 0.92 |
| 2MA6 | NMR | 0.68 | 0.75 | 0.72 | **0.82** | **0.99** | **0.92** | **0.96** |
|  | AF2 | **0.75** | **0.79** | **0.77** | 0.80 | **0.99** | **0.92** | 0.95 |
| 2RN7 | NMR | 0.52 | **0.69** | 0.61 | **0.80** | 0.95 | **0.85** | **0.90** |
|  | AF2 | **0.66** | 0.68 | **0.67** | 0.70 | **0.96** | 0.82 | 0.88 |

<sup>a</sup>DP<sub>min</sub> - lowest DP score across the ensemble.
<sup>b</sup>DP<sub>max</sub> - highest DP score across the ensemble,
<sup>c</sup>AF2 model with highest DP<sub>max</sub>: 2HEQ - Rank 1,3 ; 2KBN - Rank 1; 2KHD - Rank 3; 2KIW - Rank 2; 2KJR - Rank 1; 2KOB - Rank 1; 2KZV - Rank 3,5; 2MA6 - Rank 5; 2RN7 - Rank 2,5.
<sup>d</sup>For each NMR / AF2 comparison, and for each metric, the better score is indicated in bold font.

As ^15^N-^1^H RDC data for two alignment media, polyethylene glycol (PEG) and stretched polyacrylamide gels (PAG), were deposited in the PDB for target PDB ID 2KZV, we also compared these observed data with RDC values calculated for the corresponding AF2 and NMR models. These results, generated with *PDBStat*, are shown in Figure 3, and the corresponding linear correlation coefficients (r^2^) and RDC Q scores are summarized in Table 5. In both media, both the NMR and AF2 models are a reasonably good fit to these RDC data; with linear correlation coefficients r^2^ of 0.90 and 0.81 for AF2 Model 1, and 0.95 and 0.86 for NMR medoid structures, in PAG alignment media and PEG alignment media, respectively. Similarly RDC Q1 scores are 0.19 *vs*. 0.26 (PAG), and 0.35 *vs*. 0.40 (PEG) for NMR medoid conformer and AF2 Model, respectively. Q2 scores show similar trends (Table 5). The similarity of these scores between NMR and AF2 models is remarkable considering that the NMR structures were in fact determined using these RDC data. However, the NMR structures are indeed a better fit to the RDC data, suggesting again that further improvement of the AF2 model accuracy is possible.

**Table 5.** Linear correlations (r^2^) and Q scores for experimental vs calculated ^15^N-^1^H RDCs for target 2KZV.

| **Model** | **Alignment Medium** | **$r^2$** | **Q1** | **Q2** |
| --- | --- | --- | --- | --- |
| NMR | PAG | 0.95 | 0.19 | 0.14 |
| AF2 | PAG | 0.90 | 0.26 | 0.21 |
| NMR | PEG | 0.86 | 0.35 | 0.25 |
| AF2 | PEG | 0.81 | 0.40 | 0.36 |

**Figure 3.**
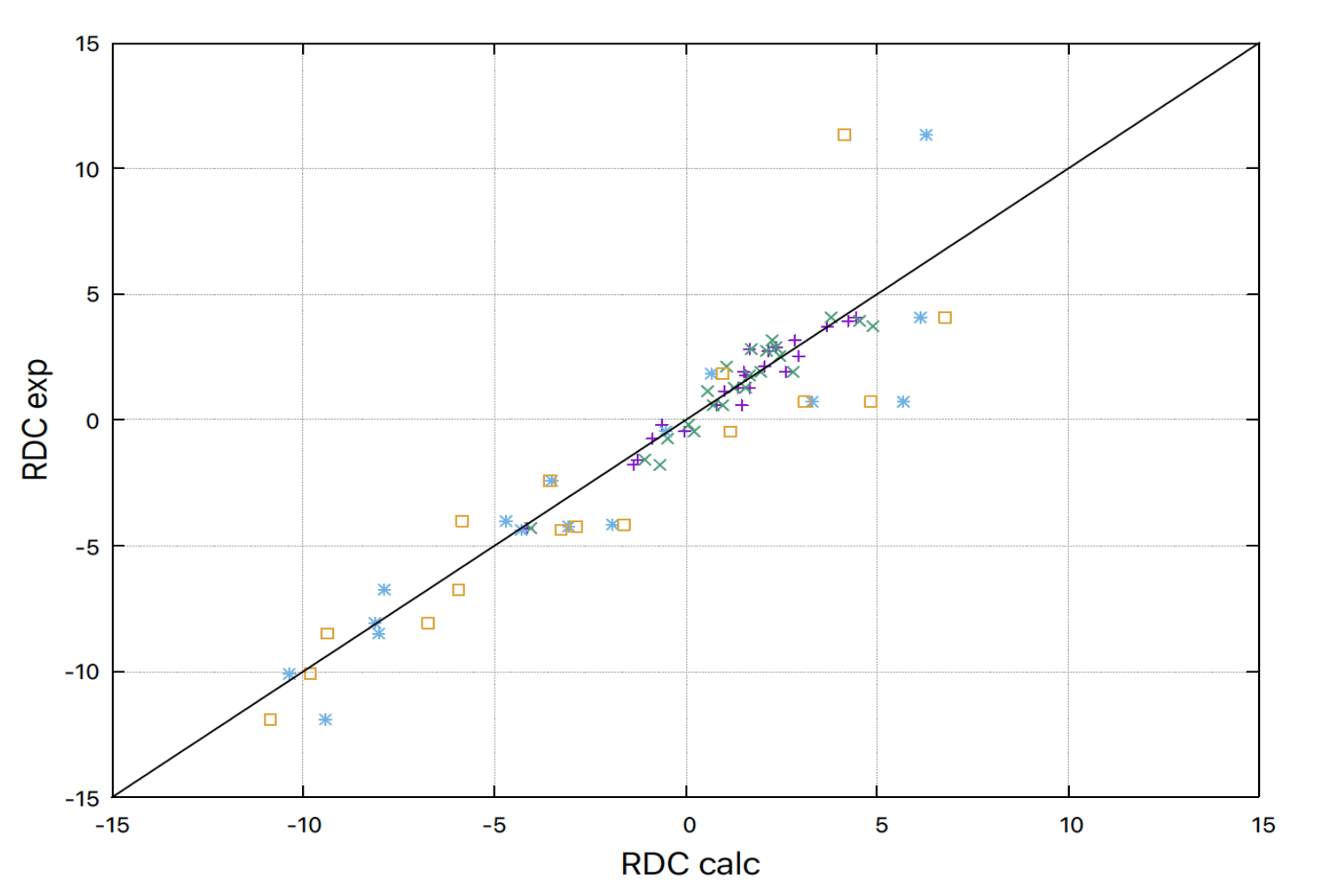
RDC analysis for target PDB id 2KZV. Experimentally-measured (RDC exp) vs calculated (RDCcalc) ^15^N-^1^H RDCs for NMR medoid conformer, PAG (violet +); NMR medoid conformer, PEG (blue *); AF2 Model 1, PAG (green X); and AF2 Model 1, PEG (orange box). Linear correlation coefficients r^2^ for these four plots are presented in Table 5.

## 4. Discussion

### AF-NMR studies

In this study, we identified 9 solution NMR protein structures which were not used in AF2 training (defined here as “blind” structures), and for which NOESY FID and peak list data are available in the public BMRB archive. Although additional data sets from the NESG and other groups may be useful for extending such analysis, these nine are sufficient to document our principal conclusion that for small, monomeric proteins not used in the training of AF2 and lacking any structural templates, AF2 models generated with the publicly-accessible AF2 CoLab server [26] have conformations that fit well to experimental data, and are very similar to models generated from extensive experimental analysis of NMR resonance assignments and NOESY data.

The Critical Assessment of Protein Structure Prediction, an experiment which has been in progress for more than 30 years, has traditionally made good use of experimental NMR structures for assessment of structure prediction methods. In CASP13 (in 2018), a NMR guided component of CASP explored the impact of sparse experimental or simulated NMR data, like that obtained for perdeuterated protein samples, on model prediction accuracy. In this experiment, it was observed that while most CASP prediction methods could not use such data reliably, a few methods could use such sparse NMR data to provide more accurate models (relative to blind reference structures) than conventional NMR structure determination methods [20]. More significantly, it was observed that in about half the target proteins studied, an early version of AlphaFold (AF1), using no experimental data, could predict protein structure models closer to the reference structures than any NMR structure modeling method tested that used these experimental data [20].

In CASP14 (in 2020), where AF2 had outstanding performance in predicting blind structures determined by X-ray crystallography and cryoEM [1], its performance on 2 of the 3 NMR targets was underwhelming. The AF2 models matched well to the NMR structure of CASP target T1055 (GDT = 0.90), but not to the experimental reference structures for CASP targets T1027 (GDT = 0.67) and T1029 (GDT = 0.47). Careful analysis of these results, and assessment against chemical shift, NOESY, and RDC data revealed highly instructive features of these two outlier NMR structures [8]. For CASP target T1027, the NMR models were found to fit most of the NMR NOESY and chemical shift data better than the AF2 prediction models. However, some of the NOESY data, which could not be explained by the NMR models, could be explained by the AF2 model. The distinct structural features of the NMR and AF2 structures of T1027, involving packing of helices into the core of the protein structure, are mutually exclusive, and these results suggest that this protein adopts multiple conformations in solution, where some NOE data fit best to the reported NMR structure and other data in the same spectra fit best to the AF2 model. This hypothesis was subsequently supported by nuclear relaxation studies on target T1027 [50], indicating intermediate exchange between two or more conformations in regions of the structure that differ between the NMR and AF2 structures. For the third CASP14 target, T1029, it was observed that the reported NMR reference structure was actually a poor fit to the NOESY peak list data (DP = 0.27). When this structure was re-determined by the original authors using more complete NOESY data, the NMR and AF2 models were observed to be in excellent agreement (GDT = 0.90) [8].

Since AF2 was made broadly available in the spring of 2021, several additional studies have examined the accuracy of structure predictions against structures determined by NMR methods. Zwecksetter compared AF2 models of three small proteins, GB3, DinI, and ubiquitin, generated using the Google Colab server, and reported excellent fits of the resulting models to experimental RDC data. However, these proteins have corresponding X-ray crystal structures that may have been used in the training of AF2. It was also not clear in this study if homologous proteins were excluded as input structural templates. Robertson, Bax and co-workers compared AF2 models against experimental RDC and NOESY data for the SARS CoV-2 main protease (M^pro^), and observed excellent concordance [51]. In this case, the analysis was done both with and without structural homologs as templates; however as there were many X-ray crystal structures of corona virus M^pro^ homologs in the PDB at the time of AF2 training, the possibility that these specific training structures significantly impacted AF2 performance on SARS CoV-2 M^pro^ cannot be excluded. Tejero et al, compared AF2 models, NMR structures, NOESY data, and RDC data for 6 proteins which had been solved by both NMR and X-ray crystallography methods [7]. These AF2 models, generated without homologous structure templates, had excellent agreement with both the NMR and X-ray structures, and also with the NMR data. They also generally had better agreement with the solution NMR structures than the corresponding X-ray crystal structures. However, the possibility that these X-ray crystal structures were used in AF2 training, and hence bias the modeling of the corresponding NMR structures, again cannot be excluded. AF2 models have also been extensively compared with NMR structures using the protein NMR structure assessment tool ANSURR [52]; this analysis included both targets with homologous X-ray crystal structures used in AF2 training, and others which may in fact be blind targets for which no homologs were available for training. However, the modeling performance on these two classes of targets was not compared. While for certain applications it is certainly appropriate to apply AF2 or other deep learning methods to model homologs (and complexes) of proteins used to train the AI and/or to use homologous protein structures as input templates for such modeling, in this study we set out to identify “blind” NMR data sets for protein domain families not used in AF2 training, and to assess prediction performance against these NMR data rather than against the atomic coordinates generated by conventional modeling methods.

### Distance restraint violations

*PSVS* uses *PDBStat* to provide an extensive and rigorous model vs distance restraint analysis [29]. NOE-based distance restraints, however, are software-(or user-) dependent interpretations of the NOESY data, involving a process for assigning NOESY cross peaks to specific interproton interactions, and calibration of interproton distances based on certain assumptions. Due to spectral degeneracy, an apparently single NOESY cross peak may arise from multiple interproton interactions, which may not be properly accounted for in creating restraints. Violated restraints are sometimes even deleted by certain software in the structure analysis process. RPF-DP scores consider all possible assignments for each NOESY cross peak and are less subjective than restraint violation analysis. For these reasons, the restraint violation analyses of AF2 vs NMR models are viewed as unreliable metrics of protein structure model accuracy and are not included in this study.

### Concordance and outliers

For 8 blind targets, the GDT between NMR (medoid) and AF2 (Model 1) models range from 0.97 - 0.92 (backbone RMSD 0.65 - 1.08 Å) and the DP scores indicate good agreement of both backbone and buried sidechain positions with the NMR data. For target PDB ID 2KIW the corresponding comparison has GDT = 0.81 (backbone rmsd 1.90 Å) (Table 2); the poorest agreement between NMR and AF2 structure models. This outlier is a 103-residue four-helical bundle domain from the DNA-binding integrase of *Staphylococcus haemolyticus*. The reported NMR structure has significantly better *Procheck-bb, Procheck-all*, and Ramachandran statistics compared to the corresponding AF2 model. The global NMR DP scores do not distinguish which model set fits the NOESY data better; the AF2 model has better DP_avg_ (0.61) and the NMR structure has better <DP> (0.74). The NMR structure 2KIW has significant differences between DP_avg_ (0.57) and <DP> (0.74), and between DP_min_ (0.50) and DP_max_ (0.61) (Table 4), which can arise from conformational dynamics, as discussed below.

### Comparing NMR and AF2 models against RDC data

Although RDC data are a rigorous method for assessing structure models, there are challenges in interpreting RDC Q scores for models that have been refined against these data [53]. Even incorrect structures may have apparently good RDC Q scores; for example the CASP14 target T1029 refined against extensive ^15^N-^1^H, ^13^Cα-C’, and Cα-Hα RDC data (in a single alignment medium) has a reasonably good Q score for these same RDC data. However, redetermination of the T1029 structure using more complete NOESY data and these same RDC data resulted in significantly different structures with excellent agreement with the more complete NOE data, and even better RDC Q scores [8]. For this protein, the significant inaccuracies of the original structure were not evident from RDC Q scores for models that were refined against the RDC data, suggesting the need for some cross-validation when using RDCs for experimental structure determination [53]. However, RDC assessments are not biased in this way when they are used to evaluate prediction models, which are generated without any sample-specific experimental RDC data. The observation that the Q scores for the NMR structure of target 2KZV, which was refined against these RDC data, are only 10 - 30% better than the models generated with AF2 demonstrate the remarkably good accuracy of the AF2 modeling of this protein structure, as well as the potential to improve the model prediction accuracy.

### Representation of solution NMR structures

Solution NMR structures are generally deposited in the PDB as an ensemble of conformations. In most cases, this ensemble is not meant to represent the distribution of conformations actually present in the sample, but rather to provide information about the convergence of the structure modeling calculations. Each conformer is considered an equally good fit to the data, within the uncertainty of the data. Accordingly, the locations of well-defined and not-well-defined regions of the model can be assessed from this ensemble [40]. Regrettably, although this concept has been an aspect of NMR structure representation since the technology for determining structures from NMR data was first developed in the mid 1980’s [46, 54-57], many users of NMR structures do not appreciate the significance of the well-defined vs not-well-defined regions of the model as they are not specifically annotated in the PDB coordinate file. AF2 provides this information in a color-encoded residue-by-residue estimate of predicted model reliability - the pLDDT score [2, 6]. Consideration of these aspects of NMR derived model reliability are essential for comparing NMR structures against prediction models, and for using NMR structures in biological applications.

### Ground truth data for assessment of improved structure prediction methods

In the most recent round of CASP15, several modeling methods were observed to outperform the standard AF2 modeling servers. All of these top-performing methods used AF2 models as input for subsequent “refinement”. As this field evolves, it is important to develop benchmark experimental data sets to assess improved methods. In particular, assessment against data (rather than against structural coordinates) has the advantage of avoiding bias introduced by the various methods used in interpreting these data as molecular models. Of special interest are solution and solid state NMR data sets, like the ones described here, for proteins not used in training these methods. For this reason, we took care to collect in a single GitHub site not only the atomic coordinates, resonance assignments, NOESY peak lists, and RDC data for these proteins, but also the corresponding NOESY and triple-resonance NMR FID data sets and PSVS / RPF structure analysis reports. This data archive, which will grow as additional validation studies like the ones outlined here are completed, should be valuable to the broader community for assessing and improving new methods of protein structure prediction and structure-guided NMR data interpretation.

### Multiple conformational states

Recently, there has been some progress using AF2 and other deep learning (DL) methods for modeling multiple native conformational states of proteins [58-60]. Ground truth NMR data for assessing such methods is especially important to develop. In this work, we report two kinds of averages for the DP score across the NMR ensemble, <DP> vs DP_avg_. The former metric treats the whole ensemble as the representation of the solution conformation, and it considers if the average interproton distances across this ensemble are consistent with the NMR NOESY data. The latter metric looks at how well each conformer of the ensemble fits the NOESY peak list data, and averages these individual fitness scores. For very tightly-bundled ensembles, these scores become identical, while for less converged ensembles they can differ by as much as 30%. Consistently, <DP> is greater than DP_avg_. Under certain circumstances, looser bundles (poorer convergence) results from dynamic averaging, in which some members of the ensemble fit a portion of the NOESY data, and other members fit a distinct portion of the data, allowing the potential for modeling multiple conformations in dynamic equilibrium from a single NOESY data set, as has been described elsewhere [8, 61, 62]. In these earlier studies, NOE data were observed to fit to a mixture of conformers predicted by computational modeling methods. In the set of nine protein targets discussed in this paper, four exhibit significant values of ΔDP = <DP> - DP_avg_ suggestive of such conformational averaging; *viz* target PDB IDs 2KHD, ΔDP = 0.12; 2KIW, ΔDP = 0.17; 2KZV, ΔDP = 0.14; and 2RN7, ΔDP = 0.19 (Table 4). In each of these cases, AF2 models fit to a subset of the NOESY data, consistent with the presence of multiple conformational states that include the AF2 structures. A more rigorous assessment of the significance of these different DP averages in terms of conformational flexibility and multiple conformational states is currently under investigation.

## 5. Conclusions

In this study, we observed that the AF2 models for nine “blind” NMR protein structures, not used in training of AF2, have accuracies, assessed by comparison against NMR NOESY, chemical shift data, and (where available) RDC data, similar to the experimental NMR models deposited in the PDB. The NOESY FID data, NOESY peak list data, chemical shifts, and model coordinates for these targets have been organized on a public GitHub site, and are accessible for assessing improved NMR data analysis and structure prediction methods. These results document the potential to use AF2 as a guiding tool to analyze NMR data and more generally for hypothesis generation in biology research.

## Abbreviations

AF: AlphaFold;
AF2: AlphaFold2;
AI: artificial intelligence;
BMRB: BioMagResDataBank;
CASP: Critical Assessment of Protein Structure Prediction;
DL: Deep Learning;
FID: Free Induction Decay data;
GDT: Global Distance Test;
NESG: Northeast Structural Genomics Consortium;
NOE: nuclear Overhauser effect;
NOESY: NOE spectroscopy;
PAG: PolyAcrylamide Gel (stretched);
PDB: Protein Data Bank;
PEG: Polyethylene Glycol;
pLDDT: predicted Local Difference Distance Test;
PSVS: Protein Structure Validation Software suite;
RDC: Residual Dipolar Coupling;
RMSD: Root Mean Square Deviation;
RPF-DP score: Recall, Precision, F-measure, and Discrimination Power score.

## Author Contributions

GTM, RT, YJH, and TAR conceived this study. EHL, LS, RT, YJH, TAR, JHP, MAK, and GTM analyzed and interpreted data. TAR, JHP, MAK, and GTM determined solution NMR structures. YJH wrote computer codes. RT wrote computer codes and carried out RDC analyses. EHL generated graphics. EHL, KF organized the GitHub Data Repository for this paper. All authors contributed in writing and editing the manuscript.

## Acknowledgements

The authors congratulate Prof. Gerhard Wagner on the occasion of his 75th birthday. GW is an inspiration to us all, and a role model of benevolent intellectual leadership for the entire biomolecular NMR community. We thank all of the scientists of the NESG consortium who produced samples, determined X-ray crystal and NMR structures, and provided these to the community by deposition in the Protein Data Bank and BioMagResDataBank. We also thank Mr. Sean Collen for his efforts in supporting our Deep Learning projects at RPI. This research was supported by grant R35-GM141818 (to GTM) from the National Institutes of Health.

## Supplementary Material

1. Supplementary Appendix A: PDBStat scripts for structural irregularity and residual dipolar coupling (RDC) analyses, and tutorial for RDC analysis with PDBStat.
2. Supplementary Table S1. Well-defined Residue Ranges for NMR and AF2 model ensembles determined by various algorithms.
3. Public GitHub site of experimental NMR data, NMR and AF2 model coordinates, and structure quality assessment reports for the 9 targets used in this study: https://github.rpi.edu/RPIBioinformatics/BlindAssessmentMonomericAF2Data

## Supplementary Appendix A

### PDBStat Scripts for Data Analysis

#### Structure Irregularity Analysis

This analysis runs the routines OMEGA, which identifies unusual trans or cis peptide bond dihedral angles, defined as values outside the ranges of 180 ± 40 deg (trans) or 0 ± 40 deg (cis) or, CIS which identifies cis peptide bonds, and MIRROR, a Cα chirality analysis identifying if each residue in the polypeptide chain is D or L.

PDBStat Script:

load pdb [file]

check

quit

#### Residual Dipolar Coupling analysis using PDBStat

a. Example data and scripts are provided at: https://github.rpi.edu/RPIBioinformatics/BlindAssessmentMonomericAF2Data/blob/main/2KZV/2KZV_realexample.tgz/)
b. Prepare input files (atomic coordinate files and rdc data files rdc.media1, rdc.media2) in format of 2KZV_realexample.tgz file folder. Ensure the residue numbering is the same for the coordinate files and RDC data files
c. Run “sh Do_2KZV” script, from 2KZV_realexample.tgz, in the directory where coordinate and RDC files are. This will create and hold the scripts for *PDBStat* and *gnuplot*, and generate the required files for RDC analysis with *PDBStat*.
d. Output files for RDC analysis will be located in the same directory, containing scripts, logs, RDC values, RDC type, and plots.

i. **Example (2kzv_RDC_all_media1/2):** **THR A 14 -1.793 -1.39 -0.69 NH** First three columns describe the residue: name, chain and number. Fourth column is the experimental RDC value. Next columns (two in this case -NMR and AF2) are the calculated RDC values for the models utilized as input. Final column is the RDC value type
ii. Q-scores Q1 and Q2 are generated in a log file
iii. Plots of RDC_exp vs RDC_calc (cf Figure 3 of main text) can be generated using gnuplot or Excel

**Supplementary Table S1.** Well-defined Residue Ranges for NMR and AF2 model ensembles determined by various algorithms.

| PDB ID | Method<br>(Medoid #)<sup>a</sup> | DAOP > 1.8 Å | Well-defined<br>CYRANGE | Well-defined<br>FindCore2 | Reliably<br>Modeled<br>AF pLDDT<sup>b, d</sup> | Consensus<br>Well-defined<br>Residue<br>Range | Comparison<br>Residue<br>Range<sup>c</sup> |
| --- | --- | --- | --- | --- | --- | --- | --- |
| 2HEQ | NMR<br>(6) | 11-20, 33-41, 46-68 | 11-20, 34-69 | 10-21, 23, 32-70 | – | 11-20, 33-41, 46-68 | 14-20, 33-41, 47-68 |
|  | AF2<br>(3) | 3-69, 72-73 | 11-69, 71-84 | 8-71 | 14-23, 32-42, 47-69 | 14-23, 32-42, 47-69 |  |
| 2KBN | NMR<br>(18) | 3-14, 17-29, 35-53, 58-77, 81-86, 89-97 | 3-29, 34-54, 58-98 | 2-30, 32-101 | – | 3-14, 17-29, 35-53, 58-77, 81-86, 89-97 | 5-14, 17-28, 35-53, 58-77, 81-86, 89-97 |
|  | AF2<br>(4) | 2-101, 103-107 | 5-28, 33-100 | 1-102 | 4-100 | 5-28, 33-100 |  |
| 2KHD | NMR<br>(1) | 31-38, 40-58, 62-69, 75-81, 84-100 | 30-97 | 25, 30-101 | – | 31-38, 40-58, 62-69, 75-81, 84-97 | 31-38, 42-58, 62-66, 75-81, 84-95 |
|  | AF2<br>(2) | 6-7, 9-14, 18-23, 25-27, 30-107 | 3-23, 30-95 | 26-108 | 30-100 | 30-95 |  |
| 2KIW | NMR<br>(9) | 5-17, 21-87 | 5-92 | 4-18, 20-94, 96 | – | 5-17, 21-87 | 5-15, 22-36, 38-87 |
|  | AF2<br>(4) | 2-99, 104-106 | 1-91 | 1-96 | 1-15, 22-36, 38-90 | 2-15, 22-36, 38-90 |  |
| 2KJR | NMR<br>(1) | 15-22, 27-52, 55-82, 85-94 | 14-95 | 13-95 | – | 15-22, 27-52, 55-82, 85-94 | 15-22, 32-52, 55-82, 85-94 |
|  | AF2<br>(5) | 15-24, 28-30, 32-94 | 3-23, 32-94 | 14-26, 28-95 | 14-23, 29-94 | 15-23, 32-94 |  |
| 2KOB | NMR<br>(5) | 63-157 | 63-157 | 61-161 | – | 63-157 | 63-156 |
|  | AF2<br>(2) | 62-158, 160-162 | 63-156 | 61-163 | 62-157 | 63-156 |  |
| 2KZV | NMR (7) | 8-21, 27-39, 45-48, 53-59, 63-80 | 7-81 | 7-22, 26-41, 43, 45, 51-68, 73-81 | – | 8-21, 27-39, 45, 53-59, 73-80 | 14-20, 27-38, 57-59, 73-80 |
|  | AF2<br>(1) | 4-20, 23-81 | 8-20, 26-38, 57-80 | 4-45, 47-48, 50-83 | 10-40, 53-81 | 14-20, 26-38, 57-80 |  |
| 2MA6 | NMR (7) | 10-17, 20-44, 49-59 | 10-58 | 8-60 | – | 10-17, 20-44, 49-58 | 10-17, 20-44, 49-58 |
|  | AF2<br>(1) | 4-5, 7-60 | 9-59 | 1-61 | 9-59 | 9-59 |  |
| 2RN7 | NMR<br>(17) | 8-25, 27-57 | 10-55 | 7-59 | – | 8-25, 27-55 | 8-25, 27-54 |
|  | AF2<br>(5) | 2-60, 68-70, 77-107 | 7-54 | 4-61 | 8-56 | 8-54 |  |

<sup>a</sup>Medoid of backbone conformations identified using *PDBStat*.
<sup>b</sup>pLDDT > 80 was used as the cutoff for reliably modeled residue ranges
<sup>c</sup>The Comparison Residue Range is the consensus between NMR and AF Consensus Residue Ranges.
<sup>d</sup>Comparing the medoid of the NMR ensemble with the Model 1 conformer of the AF2 ensemble (and the best score across all AF2 models)

